# Increased calcium spark frequency and variability of action potential duration precede early after depolarisations in isolated ventricular myocytes

**DOI:** 10.64898/2026.05.09.723211

**Authors:** S.J. Briston, D.A. Eisner, K.M. Dibb, L.A. Venetucci, A.W. Trafford

## Abstract

Drug-induced inhibition of the delayed rectifier potassium (*I*_Kr_) current predisposes to early afterdepolarisations (EADs) and cardiac arrhythmias. Here, we sought to determine the contribution of action potential duration (APD), APD variability and spontaneous calcium release from the sarcoplasmic reticulum (SR) in the formation of EADs. In isolated sheep ventricular myocytes, EADs were induced by combined inhibition of *I*_Kr_ with dofetilide and β-adrenergic stimulation. The onset of EADs was preceded by increased beat-to-beat variability of APD. To isolate the role of APD in EAD initiation, the sarcoplasmic reticulum (SR) was depleted of calcium with caffeine. The first beat post-caffeine was associated with prolonged APD but not an EAD. During β-AR stimulation, increasing ryanodine receptor open probability had no effect on APD but increased APD variability and induced both EADs and delayed afterdepolarisations (DADs). Targeting RyR open probability with K201 reversibly abolished afterdepolarisations. APD variability was a better predictor of EADs than APD alone. During an EAD, changes in [Ca^2+^]_i_ preceded those of membrane depolarisation and the changes in [Ca^2+^]_i_ were in the form of calcium sparks. In silico modelling demonstrated that membrane time constant effects account for the delay between changes in [Ca^2+^]_i_ and membrane potential. In summary, using a drug-induced model of action potential prolongation with β-AR stimulation, EADs are preceded by increased APD variability and an increase in Ca^2+^ sparks. Targeting SR function abolishes EADs. These results suggest a key role for SR Ca^2+^ overload in the formation of EADs and indicate that EADs and DADs share common mechanisms.

**Key Points:**

- Drugs that prolong the cardiac action potential and ECG QT interval are a major cause of early afterdepolarisations and dangerous ventricular arrhythmias initiated by early afterdepolarisations.
- Prolongation of the action potential is widely assumed to be the primary driver of these events.
- We show that early afterdepolarisations are instead preceded by increased beat-to-beat variability of action potential duration and that this variability has better sensitivity and specificity for early afterdepolarisations than action potential duration.
- Small, spontaneous calcium release events known as calcium sparks occur before membrane depolarisation driving early afterdepolarisations.
- Suppressing calcium release from the sarcoplasmic reticulum abolishes early afterdepolarisations, identifying calcium handling instability as potentially a key mechanism of drug-induced arrhythmia.

## Introduction

Torsade de Pointes (TdP) is a characteristic form of polymorphic ventricular tachycardia which can degenerate into fatal ventricular fibrillation. Early after-depolarisations (EADs) are secondary depolarisations occurring during either the plateau (phase 2) or early-repolarisation (phase 3) phases of the action potential and are linked to the onset of TdP by providing both the trigger and substrate for the arrhythmia (Volders *et al*., 2000). Here, the trigger component arises from the membrane depolarisation effect and the substrate component derives from the electrical heterogeneity introduced into the myocardium. EADs and thence TdP are usually associated with an abnormally long action potential repolarisation phase or prolonged electrocardiogram QT interval and are frequently linked to a reduction in repolarising potassium currents such as the delayed rectifier potassium current, *I*_Kr_. Such reductions in *I*_Kr_ are either congenital (Long QT syndrome type 2; cLQT2), a consequence of electrolyte imbalance or occur in response to drug-induced loss of channel function leading to an acquired form of QT prolongation (aQTp) (Ravens & Cerbai, 2008). Mechanistically, it has been argued that the prolongation of action potential duration (APD) increases the L-type Ca^2+^ channel window current causing reactivation of the L-type Ca^2+^ current (*I*_Ca-L_), a hypothesis supported by pharmacological augmentation of *I*_Ca-L_ inducing EADs (Marban *et al*., 1986; January & Riddle, 1989), inhibition of *I*_Ca-L_ abolishing EADs (Marban *et al*., 1986; Milberg *et al*., 2012) and experiments manipulating *I*_Ca-L_ window current using the dynamic clamp technique (Kettlewell *et al*., 2019).

However, the requirement for increased action potential duration, and QT prolongation and *I*_Ca-L_ reactivation is debated with some (Qu *et al*., 2013; Horvath *et al*., 2015) but not all (Priori & Corr, 1990) studies indicating increased APD causes EADs. Additionally, it has been argued that whilst APD and QT prolongation are important cofactors, the induction of EADs may also require an increased dispersion of repolarisation across the ventricular wall and instability in the beat to beat action potential duration and thence repolarisation, known as “beat-to-beat variability of repolarisation” (BVR) (Antzelevitch, 2007; Johnson *et al*., 2010; Dries *et al*., 2020; Amoni *et al*., 2022). Respectively, APD prolongation and increased BVR could lead to an increase in and instability of sarcoplasmic reticulum (SR) Ca^2+^ content and thereby predispose to the spontaneous release of Ca^2+^ from the SR (Diaz *et al*., 1997; Jiang *et al*., 2005; Kashimura *et al*., 2010).

Delayed after-depolarisations (DADs) are also known to initiate triggered activity driven arrhythmias in diverse conditions including cardiac glycoside toxicity (Ferrier *et al*., 1973; Di Gennaro *et al*., 1984) and catecholaminergic polymorphic ventricular tachycardia (CPVT) (Jiang *et al*., 2004; Kashimura *et al*., 2010). Here, SR Ca^2+^ content is critical and once a threshold SR Ca^2+^ content has been reached, spontaneous release of Ca^2+^ from the SR occurs giving rise to Ca^2+^ waves (Diaz *et al*., 1997) and Ca^2+^ efflux from the cell by the electrogenic Na^+^-Ca^2+^ exchanger (NCX) (Lederer & Tsien, 1976) resulting in membrane depolarisation (Ferrier *et al*., 1973).

Whilst it is widely held that EADs and DADs have distinct cellular mechanisms there is emerging evidence that, at least under some conditions, they may share a common dependence on SR load and NCX activation (Volders *et al*., 1997; Volders *et al*., 2000). However, considerable uncertainty remains regarding the role of spontaneous Ca^2+^ release in triggering EADs with, for example, some studies indicating that spontaneous Ca^2+^ release can drive EADs whilst others suggest that reactivation of *I*_Ca-L_ is required (Zhao *et al*., 2012; Song *et al*., 2015; Wilson *et al*., 2017). Additionally, inhibition of SR function with ryanodine, thapsigargin, K201 or the phosphodiesterase 5 inhibitor sildenafil has been shown to abolish EAD occurrence in several models indicating a key role for the SR in EAD formation (Burashnikov & Antzelevitch, 1998; Qi *et al*., 2009; Nemec *et al*., 2010; Zhao *et al*., 2012; Chang Liao *et al*., 2015; Kim *et al*., 2015; Hutchings *et al*., 2021). However, in these studies the detailed mechanisms were not fully elucidated, and several key questions remain regarding the cause of SR Ca^2+^ release and whether or not this is a driver of EADs in much the same way as it is in DADs.

Given the above considerations the purpose of this study was to determine the role of action potential duration, action potential instability and Ca^2+^ release from the SR in the initiation of early after-depolarisations. EADs were induced in sheep ventricular myocytes using a drug-induced model of aQTp with the *I*_Kr_ blocker dofetilide during non-specific β-adrenoceptor stimulation using isoprenaline thus mimicking a key aspects of cLQT2, impairment of *I*_Kr_. The major findings were: i) EAD formation required combined inhibition of *I*_Kr_ and β-adrenergic stimulation; ii) EADs occurred following an increase in BVR; iii) action potential prolongation alone is not sufficient to induce EADs; iv) Ca^2+^ sparks precede and thus appear to drive EAD occurrence and, iv) reducing RyR open probability supressed the formation of EADs. We therefore conclude that there is a key role for SR Ca release in the development of EADs during combined *I*_Kr_ blockade and β-adrenergic stimulation.

## Methods

### Ethical Approval

All procedures involving the use of animals were conducted in accordance with The UK Animals (Scientific Procedures) Act, 1986 and European Union Directive 2010/63. Local approval was obtained from The University of Manchester Animal Welfare and Ethical Review Board. The reporting of experiments involving animals follows the ARRIVE 2.0 guidelines where appropriate (Percie du Sert *et al*., 2020). Sheep were group housed in pens on a 12:12 hour light:dark cycle with *ad libitum* access to food and fresh water.

### Cell Isolation

Single left ventricular myocytes were isolated from the mid-myocardial wall of adult (∼ 18 months old) female Welsh Mountain sheep using a collagenase digestion as previously described (Dibb *et al*., 2004; Briston *et al*., 2011; Caldwell *et al*., 2014; Lawless *et al*., 2019). Briefly, sheep were killed with intravenous pentobarbitone (200 mg/kg). Heparin (10,000 units) was used to prevent intracoronary thrombosis and the heart was rapidly excised and placed in a Ca^2+^-free isolation solution containing (in mmol/L): NaCl 134, glucose 11; HEPES 10; 2,3-butanedione monoxime 10; KCl 4; MgSO_4_ 1.2; NaH_2_PO_4_ 1.2 and bovine serum albumin 0.5 mg/ml, pH 7.34 with NaOH. The ventricles were separated from the atria and the left descending coronary artery cannulated and perfused with the Ca^2+^–free solution at 37 °C for 10 mins to allow washout of blood. Collagenase type IV (Worthington, UK) and protease type XIV (Sigma-Aldrich, UK) were then perfused and the heart digested for ∼ 6 - 8 mins. A taurine-containing solution (in mmol/L); NaCl 113; taurine 50; glucose 11; HEPES 10; 2,3-butanedione monoxime 10; KCl 4; MgSO_4_ 1.2; NaH_2_PO_4_ 1.2; CaCl_2_ 0.1 and bovine serum albumin 0.5 mg/mL, pH 7.34 with NaOH was then perfused for 20 mins. The mid-myocardial layer was dissected out and the tissue finely minced and filtered through gauze to isolate single myocytes. Myocytes were stored at room temperature in a control solution containing (in mmol/L): NaCl 140; glucose 10; HEPES 10; KCl 4; CaCl_2_ 1.8; MgCl_2_ 1; probenecid 2, pH 7.34 with NaOH and used within 12 hours of cell isolation.

### Patch clamp recording and intracellular Ca^2+^ measurements

Action potentials (0.25 - 0.5 Hz) were recorded with the current clamp facility of the Axoclamp-2B amplifier (Molecular Devices, UK) using the perforated patch clamp technique. Patch pipettes (2 - 3 MΩ resistance) were filled with (in mmol/L): KCH_3_O_3_S 125; KCl 20; NaCl 10; HEPES 10; MgCl_2_ 5; pH 7.2 with KOH. Amphotericin-B (240 µg/mL) was added to the pipette solution to gain electrical access to the cell. Changes in intracellular Ca^2+^ were recorded by loading the cells for 10 mins with the acetoxymethylester forms of either Fura-4F (100 nmol/L) (Molecular Probes, UK) or Fura-2 LowAff (50 nmol/L) (TEFLABS, US). Cells were allowed to de-esterify for 30 mins prior to experimentation. Cells were perfused at 37 °C with control solution and EADs generated by adding dofetilide (5 µmol/L. Sigma-Aldrich, UK) and either 1 or 10 nmol/L isoprenaline in the perfusate. APD_90_ variability was measured as a coefficient of variability of time to 90 % action potential repolarisation for 5 consecutive beats not followed by an EAD and the 5 beats preceding an EAD.

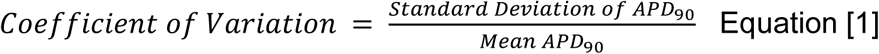

For experiments measuring NCX currents, the whole cell voltage clamp was employed using a stylised AP waveform similar in duration to that observed in cells during application of isoprenaline and dofetilide. Voltage control was achieved using an Axoclamp-2B amplifier (Molecular Devices, UK) with series resistance errors overcome using the switch clamp facility. Patch pipettes were filled with K-EGTA pipette solution (in mmol/L): KCl 120; TEA-Cl 20; HEPES 10; Na_2_ATP 5, K-EGTA 0.02, pH 7.2 with KOH. Changes in intracellular Ca^2+^ were measured with the addition of 50 µmol/L Fluo-8 (K_5_ salt; AAT Bioquest, UK) to the patch pipette. Cells were perfused at 37 °C with a control solution containing (in mmol/L): 4-aminopyridine 5; BaCl2 0.1; DIDS 0.1; Isoprenaline (10 nmol/L) and dofetilide (5 µmol/L) were used to induce EADs. For all patch clamp studies, data were digitised at 2.5 kHz.

In some experiments Ca^2+^ transients and Ca^2+^ sparks were measured by confocal microscopy (Zeiss 7-live). Cells were loaded with Fluo-8 AM (8 µM) (AAT Bioquest, UK) for 30 mins and membrane potential and action potentials measured using the perforated patch current clamp (as described above). Intracellular Ca^2+^ concentration ([Ca^2+^]_i_) was measured using a Zeiss 7Live confocal microscope (excitation 488 nmol/L, emission 525 ± 25nm) in either time-series (*xyt*; 200 Hz frame rate) or line-scan (*xt*; 1 kHz sampling rate) acquisition modes. Ca^2+^ sparks were identified using either Cardiac Spark Lab for *xt* images (Puglisi *et al*., 2014) or a custom MATLAB program for the *xyt* images.

### Membrane potential reconstruction from fluorescence signals

Membrane potential (*E_m_*) was reconstructed from the [Ca^2+^]_i_ fluorescence recordings using a passive membrane model where membrane voltage changes are determined by current flow and membrane resistance (*R_in_*) and capacitive (*C_m_*) properties. Non-linear membrane input resistance dependence on *E_m_* was derived from Powell *et al*. (1980). [Ca^2+^]_i_ fluorescence signals (*F*) were background subtracted and normalised to the resting fluorescence and used as a proxy for transmembrane current (*I_F_*) and scaled using an unconstrained scaling parameter (*g*) for each time point (*t*) (Eqn. 1).

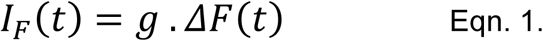

Membrane voltage changes were then described using a forward Euler approach with *R_in_* updated at each time step (*t*) based on the instantaneous *E_m_*. The fluorescence to current gain parameter (*g*) was estimated using a least squares estimation to minimise the difference between the modelled and experimentally measured voltage (Eqn. 2). An assumed membrane capacitance of 130.2 pF was used for modelling purposes (Briston *et al*., 2011).

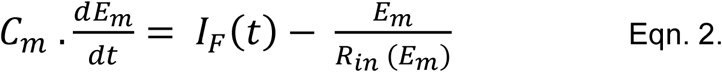

### Statistics

Data are presented as scatter plots showing all individual data points and summarised as mean ± standard deviation for *n* cells. Paired data were analysed using a Student’s t-test or repeated measures ANOVA. Data that was not normally distributed was log_10_ transformed prior to statistical testing where appropriate (Keene, 1995; Ennos, 2007) or, if normalisation was not possible a Wilcoxon signed rank test was used. Rates or proportions were analysed using Chi-squared or Fishers Exact tests. Significance was assumed when p < 0.05 and exact p values provided for p > 0.0001.

## Results

### Simultaneous I_Kr_ blockade and β-adrenergic stimulation are required to generate EADs

The first series of experiments were designed to identify the conditions required to reliably induce EADs in isolated sheep ventricular myocytes. Representative action potential traces are shown in Fig. 1A and the cumulative arrhythmia incidence summarised in Fig. 1B. During perfusion with control solution only 3.8 % of cells developed DADs and none had EADs. When *I*_Kr_ was blocked with dofetilide, action potential prolongation was observed but neither EADs nor DADs occurred (APD_90_ increasing from 360 ± 121 ms to 516 ± 139 ms; *n* = 8 cells; p *=* 0.009; Fig. 1A & C). β-adrenergic stimulation with isoprenaline had no effect on APD_90_ but was associated with the occurrence of DADs in 27.5 % of cells but EADs were not observed. Conversely, the combined inhibition of *I*_Kr_ with dofetilide and β-adrenergic stimulation with isoprenaline led to afterdepolarisations occurring in 96.2 % of cells (EADs only, 66 %; DADs only, 3.8 %; both EADs and DADs, 26.4 %). Additionally, in the 25 cells (3.8 % of total) where neither EADs nor DADs occurred during combined *I*_Kr_ blockade and β-adrenergic stimulation, a marked prolongation of APD_90_ was still observed (control, 439 ± 101 ms; Iso/Dof, 653 ± 136 ms, p < 0.0001). Additionally, compared to *I*_Kr_ blockade alone, APD_90_ was 27 ± 43 % longer in the combined presence of *I*_Kr_ blockade and β-adrenergic stimulation (p = 0.0195). Importantly, the onset of EADs and DADs during β-adrenergic stimulation or combined *I*_Kr_ blockade and β-stimulation was associated with a small (2.6 and 3.5 mV respectively) hyperpolarisation in resting membrane potential (Fig. 1D). Thus, as in other species (Jost *et al*., 2005; Xiao *et al*., 2008), sheep left ventricular myocytes have considerable repolarisation reserve and require an additional ‘hit’, in this case β-adrenergic stimulation, before EADs arise.

**Figure 1.**
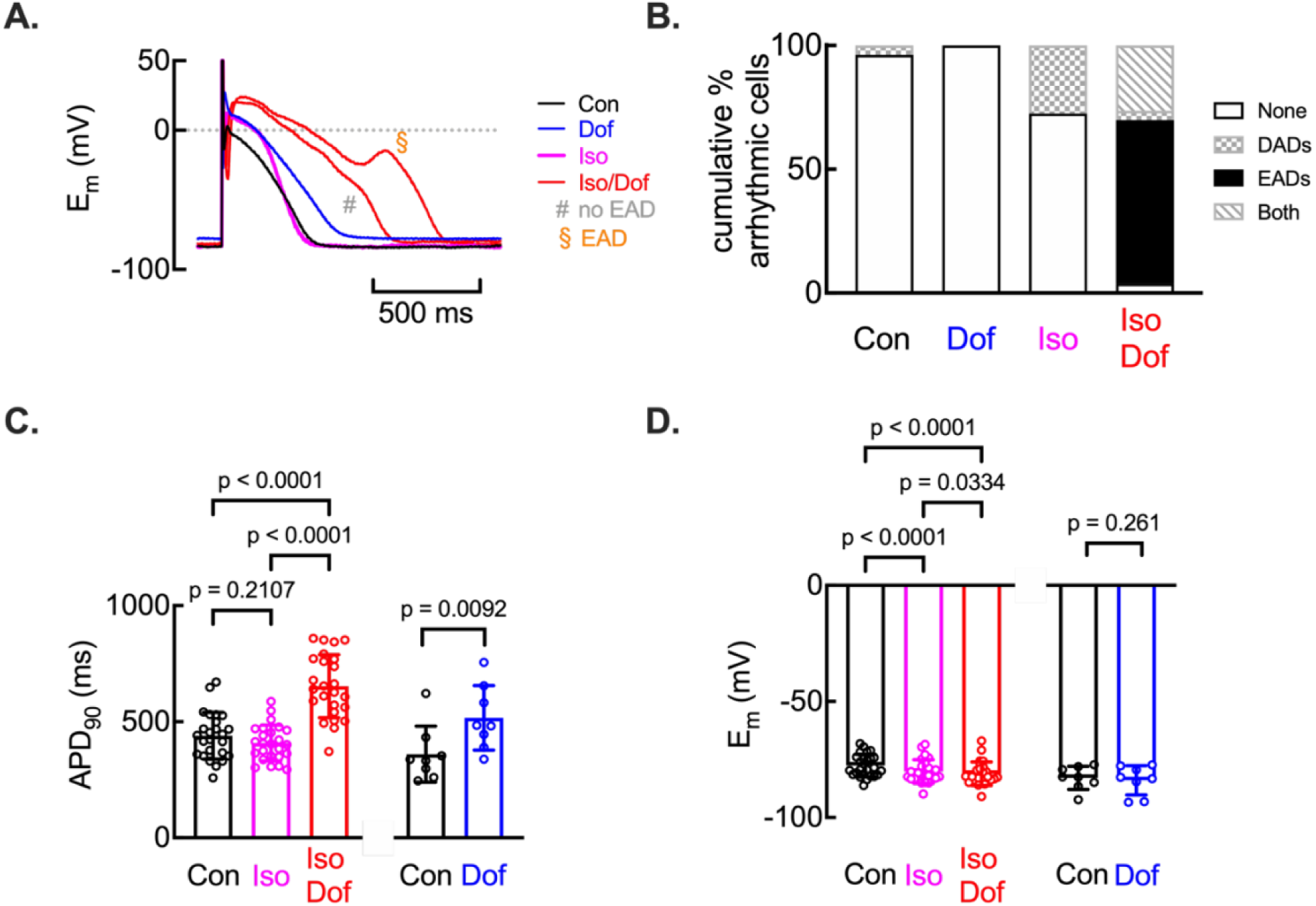
Early after depolarisations mediated by simultaneous *I*_Kr_ blockade and catecholamine stimulation in isolated ventricular myocytes. **A.** Representative action potentials under control (Con), *I*_Kr_ blockade (Dof), catecholamine stimulation (Iso) and combined *I*_Kr_ blockade and catecholamine stimulation (Iso/Dof). ^#^, action potential lacking EAD; ^§^, action potential with EAD. **B.** Summary data showing percentage of cells under control (Con), *I*_Kr_ inhibition (Dof), catecholamine stimulation (Iso) and combined *I*_Kr_ blockade and catecholamine stimulation (Iso/Dof) showing lacking EADs or DADs (None), showing DADs (DADs), EADs (EADs) or both EADs and DADs (both). **C.** Summary APD_90_ data. **D.** Summary resting membrane potential data. For **C.** and **D.** left-hand three bars are repeated measurements in same cell (One Way RM ANOVA, Tukey adjustment) and right-hand bars are paired measurements (Paired t-test, two-tailed).

### Action potential duration and beat to beat variability of APD_90_ in the initiation of EADs

Given previous studies have noted that instability in action potential duration or QT interval associates with or precedes the occurrence of afterdepolarisations or arrhythmia (Lu *et al*., 2006; Johnson *et al*., 2013; Dries *et al*., 2020; Amoni *et al*., 2022) we next sought to determine, in cells with intermittent EADs, if the onset of EADs during *I*_Kr_ blockade with dofetilide and β-adrenergic stimulation with isoprenaline was associated with changes in APD duration and augmented BVR. Figure 2A shows a representative time course of action potentials from the same cell obtained during combined *I*_Kr_ inhibition and β-adrenergic stimulation. It is clear there is variation of APD_90_ during periods of stimulation where EADs do not occur. However, beat-to-beat variation in APD_90_ nearly doubled in the five beats that preceded an action potential with an EAD (Fig. 2A & B; coefficient of variation increasing on average from 3.36 ± 1.48 % to 7.61 ± 3.39 %; n = 24 cells; p <0.0001). Additionally, in these cells APD_90_ was also longer in the five beats preceding an EAD (Fig. 2C; no EAD, 690 ± 132 ms; EAD, 808 ± 142 ms; p < 0.0001).

**Figure 2.**
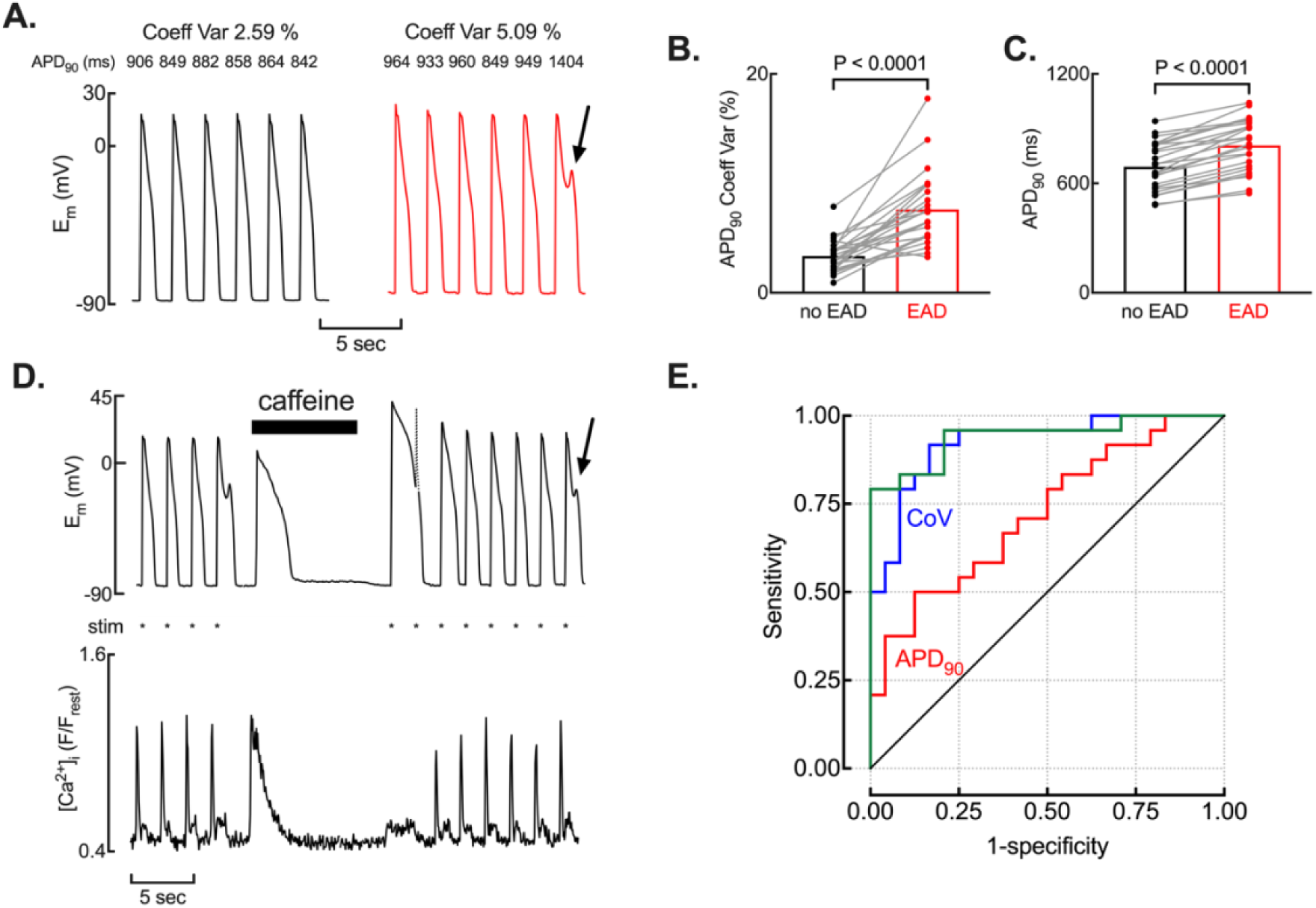
Increased beat to beat variability in action potential duration precedes early afterdepolarisations during *I*_Kr_ blockade and β-adrenergic stimulation. **A.** Representative time course showing a series of six consecutive action potentials lacking EADs (left) and, at a different time point in the same cell during *I*_Kr_ blockade with dofetilide and β-adrenergic stimulation with isoprenaline, where an EAD is observed on the sixth beat (right). Beat to beat action potential duration (APD_90_) and calculated coefficient of variation in APD_90_ are shown above. **B.** Summary data showing increased coefficient of variability in APD_90_ in beats immediately preceding EADs (Paired t-test, two-tailed). **C.** Summary data showing increase in the mean APD_90_ immediately preceding EADs (Paired t-test, two-tailed). **D.** Representative time course showing (upper) membrane potential and (lower) intracellular Ca^2+^ concentration. Caffeine (10 mmol/L) was applied as indicated and cells stimulated at 0.25 Hz (*, stimulation mark). Note the prolonged action potential on resumption of stimulation after caffeine removal (broken line, simulation artefact) and recurrence of EADs on the seventh evoked action potential (arrow). **E.** Receiver Operating Characteristic curve showing the discriminatory value of APD_90_ (red), APD_90_ variability (CoV, blue) and combined model (green).

To determine the role of action potential duration *per se* in the occurrence of EADs we used caffeine (10 mmol/L) to empty the SR of Ca^2+^ and therefore also abolish the effects of Ca^2+^ dependent inactivation of *I*_Ca-L_ on APD. Fig. 2D shows a simultaneous recording of membrane potential and [Ca^2+^]_i_ in a cell stimulated at 0.5 Hz in the presence of isoprenaline and dofetilide. When an EAD was observed, stimulation was stopped and 10 mmol/L caffeine applied to empty the SR. Following the decay of the caffeine induced Ca^2+^ transient, caffeine was removed and stimulation resumed. The duration of the first post-caffeine action potential was markedly prolonged yet no EAD was observed. On average, the first post caffeine APD_90_ was 1765 ± 732 ms, (*n =* 13 cells) and in some cells exceeded the inter-beat interval (2 seconds; e.g. Fig. 2D). Despite the profound increase in APD_90_ under these conditions of SR depletion, no EADs occurred during the first post-caffeine action potentials. However, as the SR refilled with Ca^2+^ during subsequent stimuli after caffeine removal, EADs returned indicating that the effects of caffeine are fully reversible. These observations are significant from two perspectives: i) they demonstrate that factors other than just action potential prolongation must contribute to EAD initiation and, ii) they are consistent with a role of SR Ca^2+^ content in the occurrence of EADs.

We next sought to evaluate the discriminatory value of APD_90_ and the coefficient of variability of APD_90_ for the occurrence of EADs using receiver operator characteristic (ROC) curve logistic modelling. The area under the curve (AUC) and 95 % confidence intervals (in parentheses) were; for the combined model, 0.94 (0.8731 – 1.000); for APD_90_ only, 0.7205 (0.5763 – 0.8647) and, for the coefficient of variability of APD_90_, 0.9236 (0.8487 – 0.9985). The difference in AUC for APD_90_ and coefficient of variability of APD_90_ was significant (Χ^2^ =5.9327, p = 0.0149) indicating that beat to beat variability in APD_90_ is a better predictor of EAD occurrence than APD_90_.

### The effect of increasing RyR open probability on EAD formation

Since the previous experiments indicated an important role for SR Ca^2+^ content in EAD formation we next attempted to determine if SR Ca^2+^ content was different on beats with or without an EAD. We attempted to assess SR Ca^2+^ content by applying 20 mmol/L caffeine (Briston *et al*., 2014). However, we were unable to apply caffeine rapidly enough after an EAD to convincingly discount any effects of SR refilling between the EAD and the 20 mmol/L caffeine-evoked SR Ca^2+^ release (Trafford *et al*., 1997). Given this limitation we instead sought to indirectly examine the effect of SR Ca^2+^ release in the initiation of EADs by increasing RyR opening probability (RyR P_o_) with lower concentrations (500 µmol/L) of caffeine (Trafford *et al*., 1998b). Here the purpose is to use caffeine to increase RyR open probability and lower the threshold SR Ca^2+^ content at Ca^2+^ release occurs (Trafford *et al*., 2000; Venetucci *et al*., 2007) and determine if such a manoeuvre initiates afterdepolarisations in cells where such events were previously absent.

In control solution (Con), 0 of 30 cells exhibited DADs and increasing RyR P_o_ with 500 µmol/L caffeine induced DADs in 3 of 7 cells (Fig. 3A & C, p = 0.005), However, EADs were never observed under these conditions. We therefore sought to augment SR Ca^2+^ content and repeated these experiments after β-adrenergic stimulation with isoprenaline (1 or 10 nmol/L). At the lower isoprenaline concentration, 500 µmol/L caffeine increased the occurrence of DADs (from 4.3 % to 65.2 % of cells; p < 0.001) but EADs were only observed in 1 of 23 cells). When cells were stimulated in the higher concentration of isoprenaline (10 nmol/L) DADs and EADs were sometimes observed before the addition of 500 µmol/L caffeine, however, caffeine application increased the occurrence of both EADs (Fig. 3Ab & C; from 13.3 to 60 % of cells; p = 0.021; *n* = 15 cells) and DADs (Fig. 3Ab & C; from 26.7 % to 86.7 % of cells; p = 0.005; *n* = 15 cells). Importantly, increasing RyR P_o_ with low concentrations of caffeine was not associated with any change in average APD_90_ (Fig. 3Ba). However, and consistent with the data presented above, the increase in afterdepolarisation occurrence in isoprenaline following caffeine application was associated with an increased coefficient of variability in APD_90_ (Fig. 3Bb; from 2.43 ± 1.6 % to 4.32 ± 4.1 % in 1 nmol/L isoprenaline; p = 0.018 and from 3.09 ± 1.1 % to 9.46 ± 8.4 % in 10 mmol/L isoprenaline; p < 0.0001). The dependence of arrhythmia incidence on the coefficient of variability of APD_90_ is examined in Fig. 3D and shows that across the conditions used here, these are highly correlated (p *=* 0.0053).

**Figure 3.**
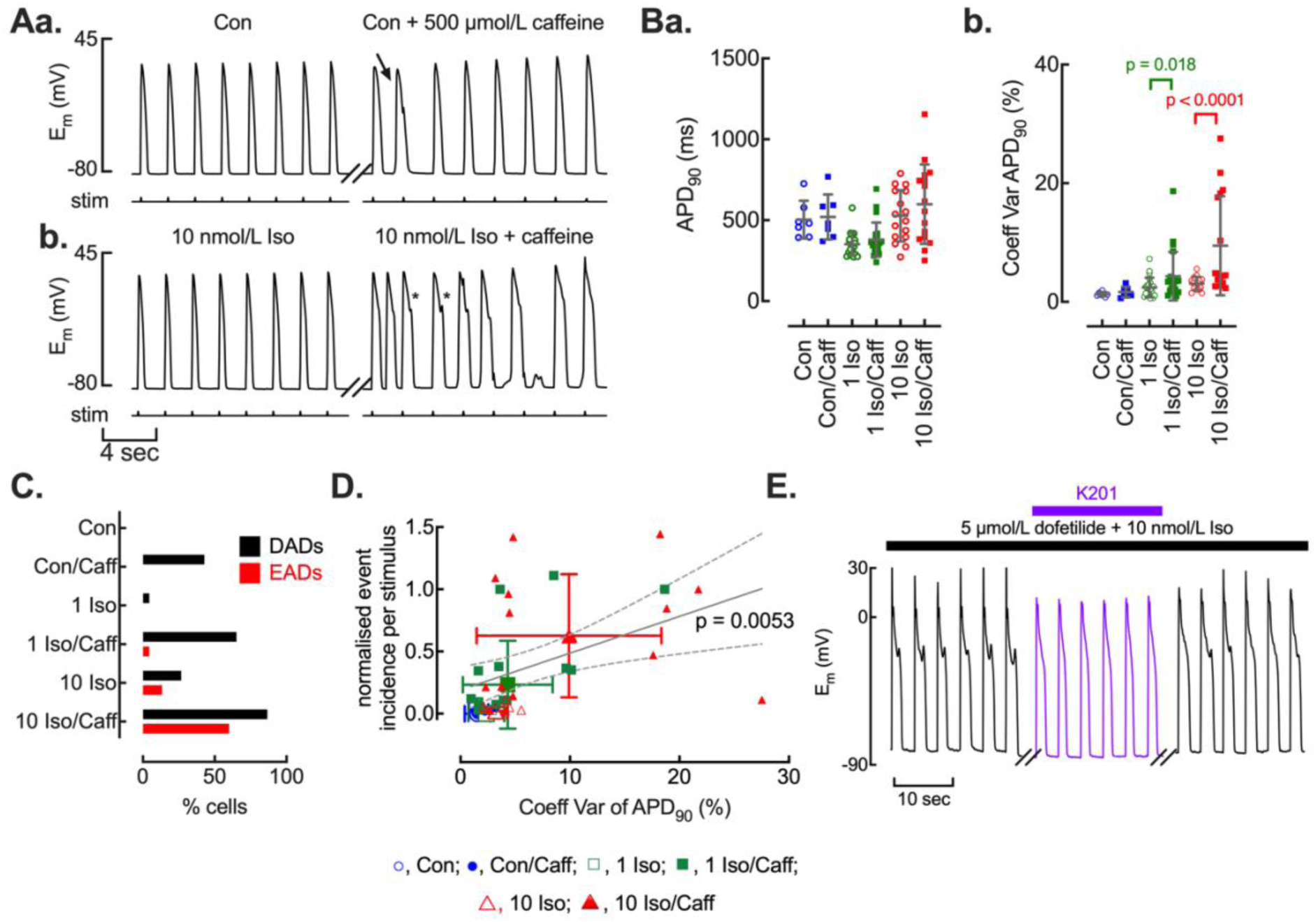
Increasing ryanodine receptor open probability augments cellular afterdepolarisations. **A.** Representative time courses showing the effect of caffeine (500 µmol/L) and catecholamine stimulation (Iso) on evoked action potentials (stim) and early and delayed after depolarisations. Caffeine addition in control conditions (Con) induced delayed afterdepolarisations (**Aa.** arrow) whereas caffeine addition during catecholamine stimulation induced both early (*) and delayed afterdepolarisation (**Ab.**). **B.** Summary data showing action potential duration (**Ba.)** and coefficient of variability of action potential duration (**Βb.**). **C.** Summary data for incidence of early (EAD) and delayed (DAD) afterdepolarisations under all experimental conditions. **D.** Relationship between arrhythmic events and the coefficient of variability of action potential duration. Large symbols denote mean data and standard deviation, small symbols show individual cell data. Linear regression (solid line) and 95 % confidence intervals (broken lines) fitted to individual data points. **E.** Experimental time course showing reversible inhibition of early after depolarisations by K201.

From these experiments it is clear that increasing RyR P_o_ with low concentrations of caffeine increases the probability of afterdepolarisations occurring. Therefore, we next sought to confirm the role of RyR P_o_ in the initiation of EADs by investigating the effect of reducing RyR P_o_ on EADs. Fig. 3E demonstrates that application of the RyR inhibitor K201 leads to a reversible abolition of EADs (in 3 of 3 experiments) in the presence of combined *I*_Kr_ blockade and β-adrenergic stimulation.

### Changes in [Ca^2+^]_i_ precede changes in membrane potential during early afterdepolarisations

The previous experiments indicate that SR Ca^2+^ content and RyR P_o_ are important cofactors in the generation of afterdepolarisations in ventricular myocytes and that early and delayed afterdepolarisations share common underpinning mechanisms. In the next series of experiments, we therefore sought to determine the role of [Ca^2+^]_i_ and membrane depolarisation. If, EADs and DADs share common mechanisms as our data suggests, then one would expect to see intracellular Ca^2+^ levels changing prior to any change in membrane potential. This is examined in detail in Fig. 4. From the representative action potential and associated Ca^2+^ transient (Fig. 4A), the secondary membrane depolarisation due to an EAD is also associated with a secondary increase in [Ca^2+^]_i_. Closer examination reveals that the secondary changes in [Ca^2+^]_i_ precede those of membrane depolarisation (Fig. 4B). On average (*n* = 9 cells) the secondary change in [Ca^2+^]_i_ occurs 20 ± 7.5 ms before those of membrane potential resulting in a hysteresis between [Ca^2+^]_i_ and membrane potential (Fig. 4C) supporting the suggestion that changes in [Ca^2+^]_i_ do drive EAD formation.

**Figure 4.**
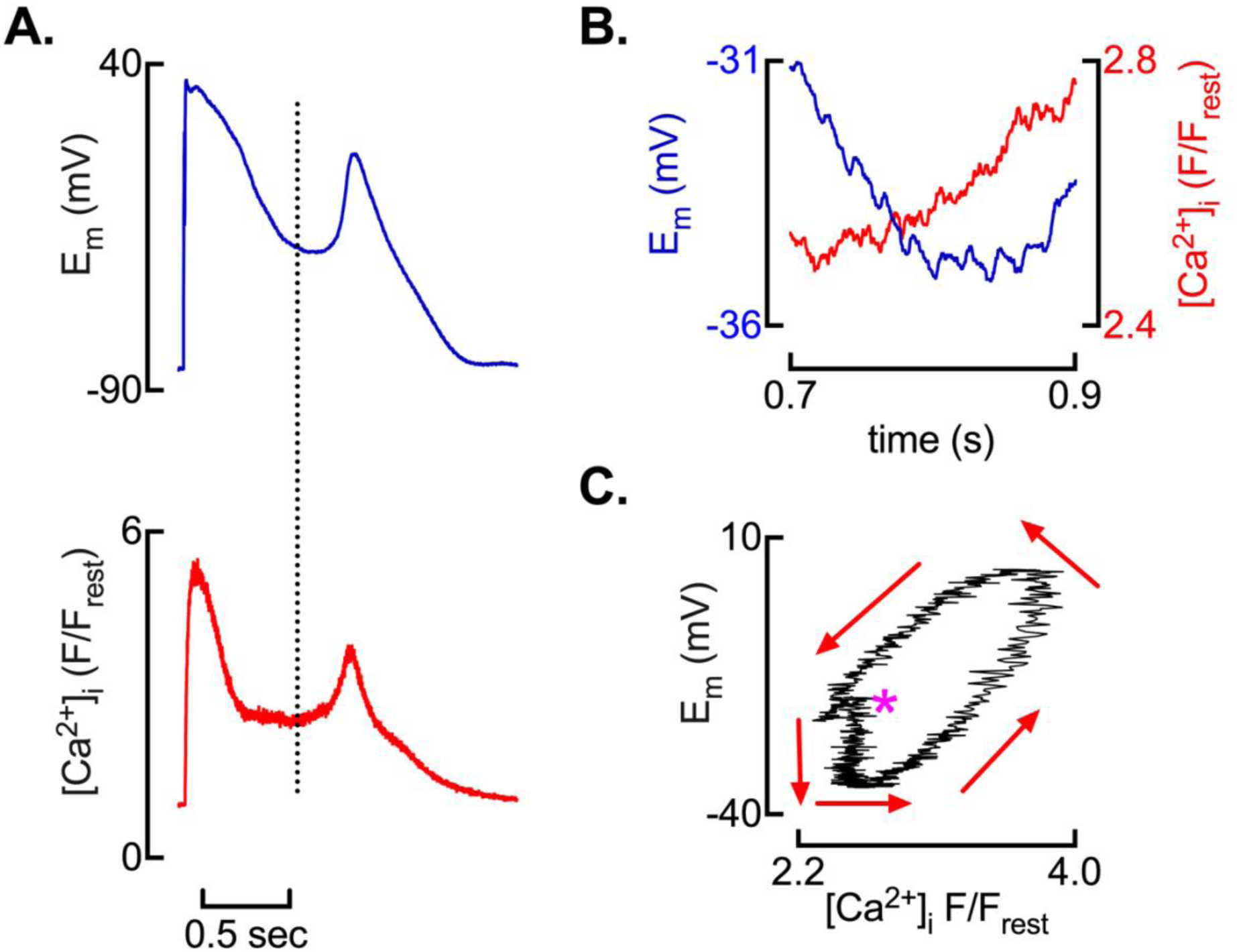
Changes in intracellular calcium concentration precede those of membrane potential during early afterdepolarisations. **A.** Simultaneous measurements of membrane potential and intracellular calcium concentration in a single myocyte showing intracellular calcium concentration increasing before membrane depolarisation (broken line). **B.** Expanded view of membrane potential and intracellular calcium changes from time shown by broken line in A. **C.** Hysteresis plot showing intracellular calcium changes before membrane depolarisation; *, denotes origin and arrows time

Given that changes in [Ca^2+^]_i_ drive Ca^2+^ dependent depolarising membrane currents such as the Na^+^-Ca^2+^ exchanger (NCX) (Lederer & Tsien, 1976), the delay between the changes in [Ca^2+^]_i_ and membrane potential during an EAD could arise as a result of membrane time constant effects arising from the product of the membrane resistance and capacitance (Time constant = Resistance * Capacitance). We took an *in silico* approach to examine this using simultaneous action potential and [Ca^2+^]_i_ measurements from the sequential EAD and non-EAD beats (Fig. 5A). Using the [Ca^2+^]_i_ difference between an EAD and non-EAD transient and the relationship between membrane potential and membrane resistance (R_in_) (Powell *et al*., 1980) as input parameters, modelling suggests the delay between changes in [Ca^2+^]_i_ and membrane potential during an EAD can be accounted for by membrane time constant effects (Fig. 5B).

**Figure 5.**
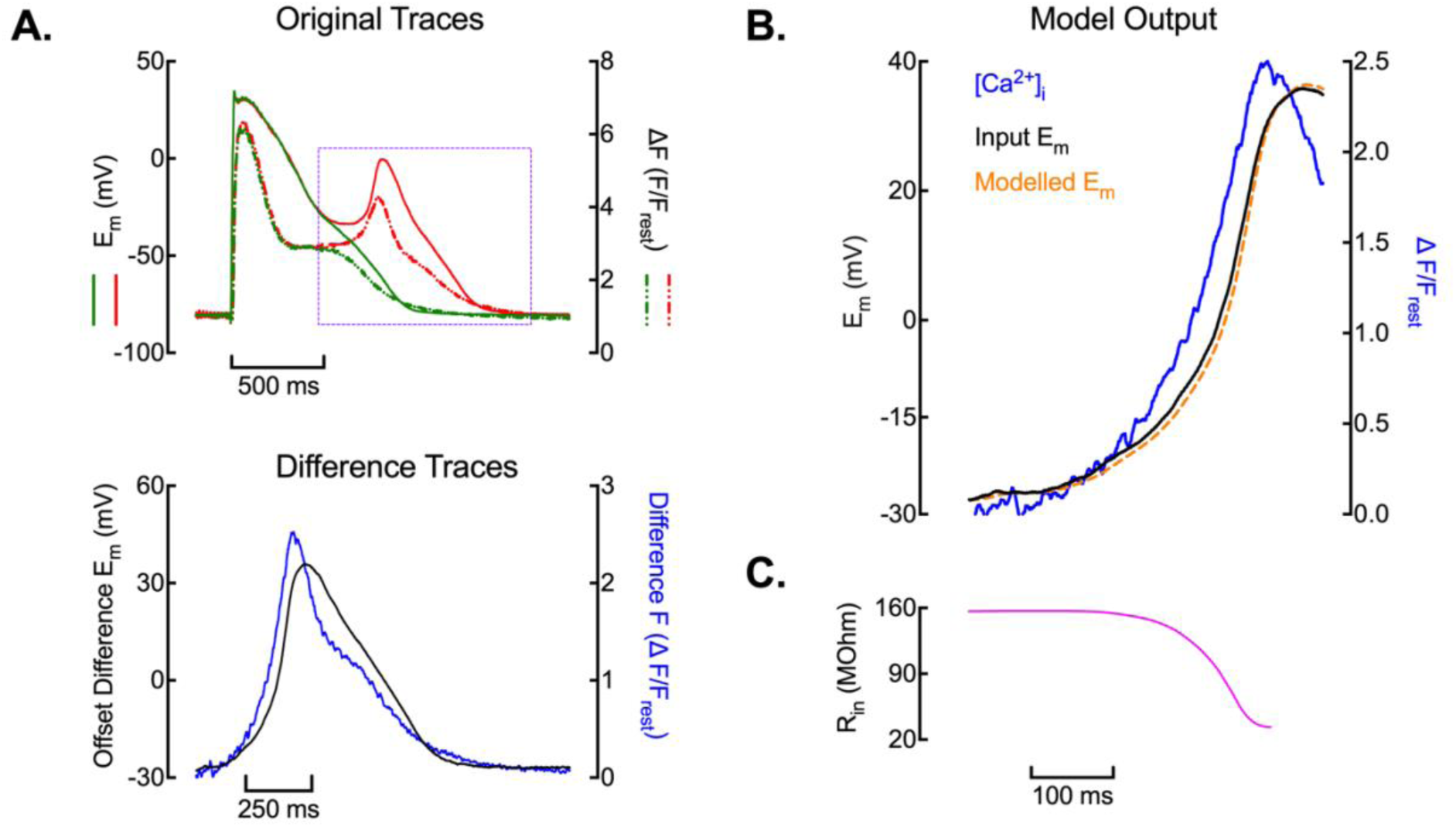
Membrane time constant effects account for delay between changes in [Ca^2+^]_i_ and membrane potential during early afterdepolarisations. **A.** Original traces (upper) from sequential transients showing membrane potential (solid line) and [Ca^2+^]_i_ (broken line) exhibiting an EAD (red traces) and without an EAD (green traces). Difference traces (lower) calculated for the region highlighted in upper panel by subtracting the no EAD trace from the EAD trace. The membrane potential difference trace has been offset vertically such that the subtracted trace originates at the actual membrane potential at the start of the period of subtraction. **B.** Modelling output focusing on the upstroke of the EAD. **C.** Input resistance changes during the upstroke of the EAD obtained from Powell *et al* (1980).

We also examined the relationship between the secondary rise of [Ca^2+^]_i_ and membrane currents in voltage clamped myocytes (Fig. 6A). During the stylised plateau phase of the depolarising pulse, secondary Ca^2+^ rises again occur prior to an inward membrane current (Fig. 6B) resulting in the hysteresis plot (Fig. 6C) where changes in [Ca^2+^]_i_ occur ahead of those of an inward, depolarising, membrane current. Additionally, consistent with our working hypothesis that changes in membrane current are Ca^2+^ dependent, the magnitude of the inward membrane current was proportional to the change in [Ca^2+^]_i_ (Fig. 6D; p = 0.003).

**Figure 6.**
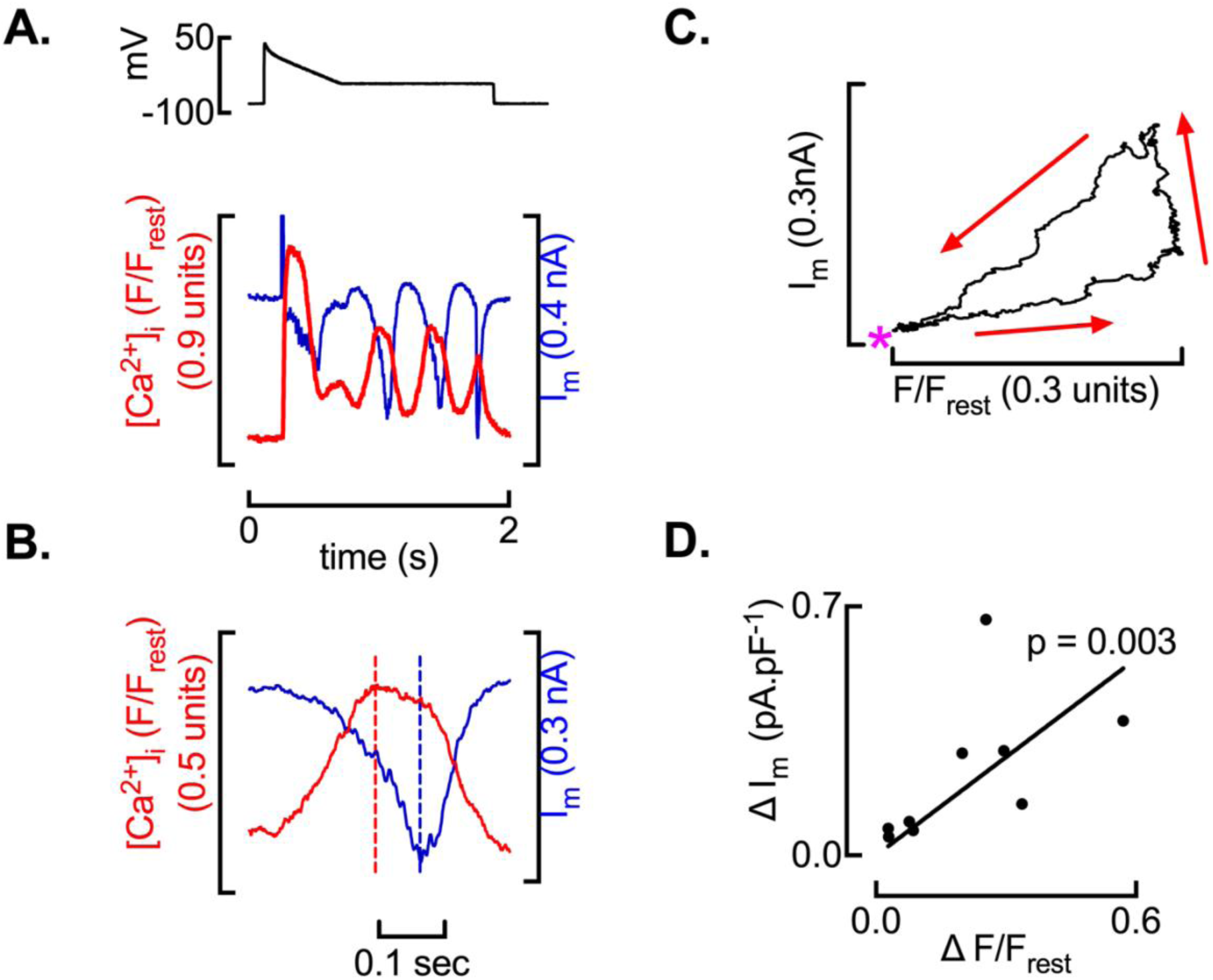
Hysteresis between [Ca^2+^]_i_ and membrane potential during secondary [Ca^2+^]_i_ elevations are maintained under voltage clamp conditions. **A.** Original traces from a voltage clamped myocyte showing voltage command waveform (upper) and changes of intracellular calcium concentration and membrane current (lower). **B.** Expanded view of changes in membrane potential and intracellular calcium concentration during a plateau phase oscillation (after depolarisation); broken lines denote peak intracellular calcium and membrane current. **C.** Hysteresis plot for the current and calcium oscillation shown in B. *, denotes origin and arrows time. **D.** Relationship between magnitude of inward current and change in intracellular calcium during plateau phase membrane oscillations. Solid line is best fit linear regression to the data with slope significantly non-zero (p = 0.003).

### EADs are preceded by an increase in Ca^2+^ sparks

The previous experiments demonstrate that changes in [Ca^2+^]_i_ drive the occurrence of EADs via generation of an inward depolarising membrane current. In the final series of experiments, we therefore sought to establish if these changes in [Ca^2+^]_i_ occurred as whole cell Ca^2+^ transients, Ca^2+^ waves or Ca^2+^ sparks. Cells were current clamped and initially imaged by high speed (200 Hz) 2-dimensional *xyt* confocal scanning. Fig. 7A shows three consecutive action potentials from a cell during combined *I*_Kr_ inhibition and β-adrenergic stimulation. In these cells, Ca^2+^ sparks are observed during the decay phase of the systolic transient. The EADs occurring on the first and third transient were associated with an increased number of Ca^2+^ sparks immediately before the afterdepolarisation (see supplementary online video). On average there was a 175 ± 111 % increase in the number of Ca^2+^ sparks when EADs occurred (Fig. 7B; no EAD, 80 ± 54 sparks/cycle; EAD, 180 ± 104 sparks/cycle; *n* = 7 cells; p *=* 0.0046).

**Figure 7.**
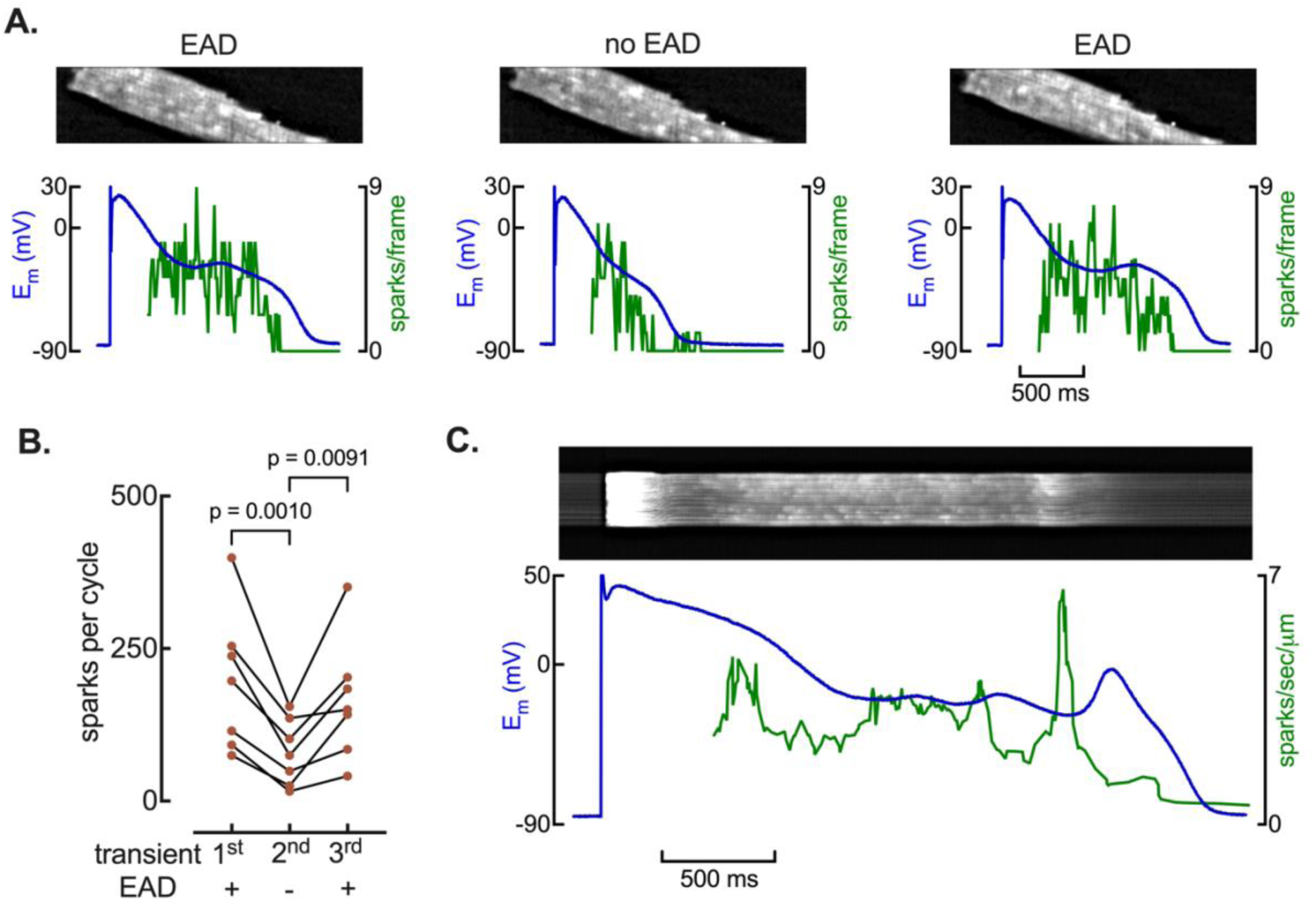
Calcium sparks mediate early afterdepolarisations. **A.** Representative plateau phase confocal (*xy* scanning, 200 Hz frame rate) images (upper) and membrane potential and calcium spark quantification (lower) from three consecutive action potentials during *I*_Kr_ blockade and catecholamine stimulation. **B.** Summary data showing plateau phase calcium spark count on consecutive action potentials in the presence (+) and absence (-) of early after depolarisations. Repeated Measures One-Way ANOVA. **C.** Representative simultaneous high temporal resolution (*xt* line scanning, 1 kHz) confocal image of intracellular calcium concentration (upper) and membrane potential and calcium spark density (lower) showing augmentation of calcium spark density preceding each early afterdepolarisation.

We also examined the relationship between Ca^2+^ sparks and EADs at higher temporal resolution using *xt* (linescan, 1 kHz) imaging (Fig. 7C). Once again it is clear that there is an increase in Ca^2+^ sparks ahead of each membrane afterdepolarisation. This pattern was observed in all (*n* = 12) imaged cells where EADs were observed. Thus, both imaging formats demonstrate that EADs elicited by *I*_Kr_ blockade and β-adrenergic stimulation occur as a result of an increase in Ca^2+^ sparks and not from whole cell Ca^2+^ transients or Ca^2+^ waves.

## Discussion

In this study we investigated the cellular mechanisms of EADs in a drug-induced model of LQT2. Our main findings were fivefold; i) *I*_Kr_ blockade and β-adrenergic stimulation were required to reliably induce EADs in isolated sheep ventricular myocytes, ii) increased variability in action potential duration preceded EADs and was a better discriminator for the occurrence of EADs than action potential duration, iii) EADs and DADs often occurred in the same cell, iv) altering RyR open probability with caffeine or K201 respectively increased and inhibited EADs and, v) an increase in Ca^2+^ sparks drives EAD formation. Taken together, these findings demonstrate an important role for beat-to-beat action potential heterogeneity and Ca^2+^ release from the SR in the formation of EADs and are consistent with EADs sharing a similar dependence on SR Ca^2+^ content and thus mechanisms of initiation to that of DADs.

### Repolarisation reserve and action potential duration

Our finding that inhibition of *I*_Kr_ alone with dofetilide failed to induce either form of afterdepolarisation despite a substantial increase in action potential duration is consistent with previous studies demonstrating considerable repolarisation reserve in the heart (reviewed in (Varro & Baczko, 2011)). Here, a decrease in one repolarising current, e.g. *I*_Kr_, is offset by a compensatory increase in other repolarising currents, e.g. *I*_Ks_ (Xiao *et al*., 2008), or reduction in depolarising currents, e.g. *I*_Na,L_ (Hegyi *et al*., 2020), to protect against the development of EADs and onset of arrhythmias. The mechanisms can involve acute counterbalancing of ion channel activity (Hegyi *et al*., 2020), transcriptional control (Xiao *et al*., 2008) and regulation of endocytic and membrane trafficking pathways (Guo *et al*., 2011). However, when a second hit is experienced, this endogenous protective repolarisation reserve effect can be overcome and arrhythmias develop (Volders *et al*., 2003; Johnson *et al*., 2013). In our acute *I*_Kr_ blockade with dofetilide experiments as a model of acquired QT prolongation, β-adrenergic stimulation provided the second hit resulting in the appearance of both early and delayed afterdepolarisations.

Whilst the addition of β-adrenergic stimulation led to both a prolongation of action potential duration and the occurrence of afterdepolarisations, two observations in our data suggest that action potential duration *per se* is a poor indicator of EADs; i) when SR function is disabled, APD_90_ is markedly prolonged, yet EADs are not observed and, ii) beat-to-beat variation in APD_90_ (CoV) has greater discriminatory value than APD_90_ for EAD occurrence. An important caveat to this conclusion is that changes in APD themselves can also influence the amount of Ca^2+^ stored within the SR (Terracciano *et al*., 1997; Mason & MacLeod, 2009). Nevertheless, our observation that when the SR is depleted following caffeine application afterdepolarisations are absent is at least consistent with factors other than solely APD driving EAD occurrence.

Numerous preclinical models (Thomsen *et al*., 2004; Johnson *et al*., 2013; Hutchings *et al*., 2021), computational (Sato *et al*., 2025a, b) and clinical studies (Hinterseer *et al*., 2008; Hinterseer *et al*., 2010; Deissler *et al*., 2026) have demonstrated increased beat-to-beat variability preceding arrhythmic events in diverse proarrhythmic settings. The variability of APD in these studies, and indeed our own ROC analysis, provides a future avenue for understanding the difference in arrhythmic risk between patients with similarly prolonged QTc intervals (Mazzanti *et al*., 2018). Our data is at least consistent with a role for heterogeneity in APD_90_ both within individual cells between beats and across different regions of the heart leading to the formation steep gradients of repolarisation that subsequently act as areas of arrhythmia initiation and / or reentry (Boulaksil *et al*., 2011; Vijayakumar *et al*., 2014; Dunnink *et al*., 2017). However, future work is required to provide a detailed characterisation of the precise ion channel, calcium regulatory or signalling mechanisms that give rise to beat-to-beat variation in APD_90_ or QTc and thence repolarisation gradient heterogeneity and arrhythmia initiation at both the cellular and intact tissue levels.

### Sarcoplasmic reticulum and ryanodine receptor function

As noted above, when we disabled SR function with a high concentration of caffeine this led to a marked prolongation of APD_90_ likely due to the loss of calcium dependent inactivation of the L-type Ca^2+^ current (Trafford *et al*., 1997). Despite the longer APD_90_, no EADs were observed on the first beat following resumption of stimulation after caffeine removal when the SR was depleted of Ca^2+^. The abolition of EADs when SR function has been disabled with either thapsigargin or ryanodine has been noted previously (Qi *et al*., 2009). However, the experiments of Qi *et al*., (2009) were performed under steady-state conditions and action potential duration was shorter when the SR was disabled, an effect which itself would also potentially contribute to a reduced likelihood of EADs if APD were to be a driver of EADs as we have shown APD_90_ is longer in cells when EADs occur.

Following the removal of caffeine and resumption of stimulation, the SR refills with Ca^2+^ (Trafford *et al*., 1997; Trafford *et al*., 1998b). The reappearance of EADs after several beats indicates an important role for SR Ca^2+^ and that EADs and DADs may share a similar dependence on a threshold SR Ca^2+^ content for their initiation (Diaz *et al*., 1997; Jiang *et al*., 2004). Additionally, β-adrenergic stimulation also increases SR Ca^2+^ content (Hussain & Orchard, 1997; Briston *et al*., 2014) and this is known to cause DADs in the inherited arrhythmia syndrome CPVT (Kashimura *et al*., 2010). Importantly, in our experiments during *I*_Kr_ blockade with dofetilide, β-adrenergic stimulation led to the occurrence of both EADs and DADs, frequently occurring together in the same cell. Despite not being able to quantify SR Ca^2+^ content immediately after an action with sufficient temporal fidelity due to the need to switch between current and voltage clamp control, we have explored the role of SR Ca^2+^ content indirectly by making use of low concentrations of caffeine to increase RyR open probability. Previously we have shown that increasing RyR open probability reduces the threshold SR Ca^2+^ content for DADs (Trafford *et al*., 2000; Venetucci *et al*., 2007). In the current experiments, both DADs and EADs were more likely when RyR open probability was increased, and particularly in the presence of β-adrenergic stimulation. Further supporting a role for increased beat-to-beat variation of action potential duration in the initiation of EADs, enhancing RyR open probability during β-adrenergic stimulation also increased the coefficient of variation in APD_90_ but this was not associated with a change in APD_90_.

The abolition of EADs by reduction of RyR open probability with K201 is also consistent with a role for Ca^2+^ release from the SR in the formation of EADs. Similar effects have been noted where K201 (JTV519) abolished spontaneous Ca^2+^ release events in expression systems (Hunt *et al*., 2007), isolated myocytes (Loughrey *et al*., 2007; Kim *et al*., 2015) and optically mapped Langendorff perfused hearts (Kim *et al*., 2013). However, in a canine *in vivo* model of dofetilide induced aQTP, K201 had no effect on arrhythmias, and at higher doses was found to increase the arrhythmias (Stams *et al*., 2011). There are at least two possible explanations for the lack of antiarrhythmic effect, or indeed the proarrhythmic effect, noted *in vivo* by Stams *et al* (2011). Firstly, in the *in vivo* experiments the higher plasma concentrations of K201 could be causing known off target effects of K201 on a number of membrane currents including *I*_Kr_, *I*_Ca-L_, *I*_Na_ and, *I*_K1_ (Kaneko *et al*., 2009). The future development of highly specific reversible blockers of the cardiac RyR would allow some of these differences to be resolved. Secondly, as we have shown extensively before (Trafford *et al*., 1998b; Choi *et al*., 2000; Trafford *et al*., 2000; Greensmith *et al*., 2014), modulation of RyR open probability only transiently affects systolic Ca^2+^ or the frequency of Ca^2+^ waves due to compensatory changes of SR Ca^2+^ content. As such, as well as targeting RyR open probability, additional strategies to prevent compensatory changes in SR Ca^2+^ content may offer more effective therapeutic pipelines. Nevertheless, the effects of low concentrations of caffeine and K201 in the present study are strongly supportive of a key role for SR Ca^2+^ release in the genesis of EADs.

### Calcium as a driver of early afterdepolarisations

The secondary depolarisations giving rise to EADs have a number of potential mechanisms including; i) reactivation or augmentation of depolarising inward currents including *I*_Ca-L_ and the late component of the sodium current (*I*_Na,L_) (Kettlewell *et al*., 2019; Ton *et al*., 2021; Alexander *et al*., 2023), ii) attenuation of repolarising or membrane stabilising currents (*I*_Kr_, *I*_Ks_, *I*_K1_) or, iii) release of Ca^2+^ from the SR (Volders *et al*., 1997; Nemec *et al*., 2010). However, these mechanisms are not mutually exclusive and under certain conditions could act in concert. For example, during hypokalaemia or hypomagnesaemia, repolarising potassium currents are impaired, prolonging action potential duration, augmenting *I*_Na,L_ leading to reverse mode Ca^2+^ entry, SR Ca^2+^ loading and thence spontaneous Ca^2+^ release from the SR and afterdepolarisations. In the present experiments, performed in normal extracellular potassium and magnesium concentrations, changes in [Ca^2+^]_i_ always preceded changes in membrane potential. Computational modelling indicates this delay can be explained through membrane time constant effects whereby changes in [Ca^2+^]_i_ lead to the instantaneous flow of a depolarising membrane current but membrane capacitance and membrane resistance effects delay the translation of this membrane current to a change in membrane potential. Similar findings, where changes in [Ca^2+^]_i_ precede membrane depolarisation have been reported before in models of drug-induced LQT2 in optically mapped hearts (Choi *et al*., 2002; Nemec *et al*., 2010; Kim *et al*., 2015).

Given that [Ca^2+^]_i_ changes drive those of membrane potential, this implies a key role for non-triggered Ca^2+^ release from the SR rather Ca^2+^-induced-Ca^2+^ release due to reactivation of *I*_Ca-L_. In optical mapping experiments the lack of spatial resolution prevents full characterisation of the nature of the Ca^2+^ release from the SR. Conversely, in the current experiments performed in single cells and using high speed *xt* or *xyt* confocal imaging, the increase in [Ca^2+^]_i_ that preceded EADs was always observed in the form of Ca^2+^ sparks occurring during the plateau phase of the action potential. A similar occurrence of late Ca^2+^ sparks and the formation of EADs has been reported in a rabbit model of heart failure (Fowler *et al*., 2020). Interestingly, in the present experiments we were able to track the relationship between Ca^2+^ sparks and the occurrence of EADs on a beat-by-beat basis. Under these conditions we noted a threshold like phenomenon between the number of Ca^2+^ sparks and the occurrence of an EAD with higher numbers of sparks being associated with the presence of EADs.

### Limitations

Whilst the findings of the present study show that Ca^2+^ sparks drive the formation of EADs and altering the open probability of the RyR modulates EAD occurrence are strongly suggestive of a role for threshold SR Ca^2+^ content in the genesis of EADs there are some methodological limitations to this study. Firstly, the time required to switch between current and voltage clamp control immediately after an EAD and change experimental solutions to block other Ca^2+^ activated currents (Trafford *et al*., 1998a; Verkerk *et al*., 2003) to isolate Na^+^-Ca^2+^ exchange current in order to quantify SR Ca^2+^ content was too long and therefore precluded assessment of the threshold SR Ca^2+^ content. Two alternative methods exist for assessing SR Ca^2+^ content; i) measuring the amplitude of the caffeine evoked Ca^2+^ transient and, ii) intra SR fluorescent dyes (Greensmith *et al*., 2014). However, both of these methods only provide a qualitative measure and are limited by the potential for non-linearity of indictor response and indicator saturation particularly under conditions of increased SR Ca^2+^ load e.g. during β-adrenergic stimulation. Secondly, our experiments have been performed on isolated myocytes and therefore any electrotonic coupling effect between cells that could dampen EAD formation has been lost. Nevertheless, in intact hearts [Ca^2+^]_i_ driven EAD formation has been noted and therefore the cellular observations in the present study closely resemble those in the intact heart. Thirdly, we used K201 and low concentrations of caffeine as tools to investigate a role for RyR mediate Ca^2+^ release as a driver of EADs. Neither of these drugs is specific for the RyR and we therefore cannot completely exclude off-target effects. However, in previous experiments we have extensively demonstrated for low concentrations of caffeine the effects are mediated through the RyR (Trafford *et al*., 1998b; Trafford *et al*., 2000). Finally, our experiments and indeed those of others, have been performed under an acute setting of *I*_Kr_ blockade as a model of aQTP. We assume our findings will be equally applicable in both congenital forms of LQT2 and where QT prolonging drugs may be given chronically. However, this and the effect that any compensatory changes in other ion channels may have will need to be evaluated in different experimental models.

## Conclusion

In summary, using a cellular model of drug-induced aQTP, we have demonstrated that *I*_Kr_ blockade and APD prolongation alone are insufficient to produce arrhythmic events and that β-adrenergic stimulation is required and induces both EADs and DADs in the same cell. Our observations that; i) EADs are driven by Ca^2+^ release from the SR and, ii) modifying RyR open probability alters EAD occurrence are strongly supportive of a shared SR dependent mechanism between DADs and EADs. In our study we were able to show that Ca^2+^ sparks drive the formation of EADs. Finally, and of particular note, we found that beat-to-beat variability of action potential duration was a much better discriminator for EADs that action potential duration. These findings are of potential significance for both the prediction of arrhythmia onset in aQTP and other congenital and acquired arrhythmia syndromes and the development of novel antiarrhythmic strategies for these conditions.

## Additional Information

### Data and code availability

On formal acceptance for publication in a peer-reviewed journal, data and coding associated with this manuscript will be made available on a Creative Commons CC-BY open access basis through The University of Manchester FigShare repository (DOI provided on acceptance)

### Author contributions

All authors have approved the final version of the manuscript.

Conceptualisation, AWT; Methodology, All authors; Investigation, SJB; Formal analysis & Validation, SJB & AWT; Writing – Original Draft, AWT & SJB; Writing – Review Editing, all authors; Visualisation, SJB & AWT; Supervision, all authors; Funding, AWT, DAE.

