## Supplemental video for "Increased calcium spark frequency and variability of action potential duration precede early after depolarisations in isolated ventricular myocytes"

### Slide 1
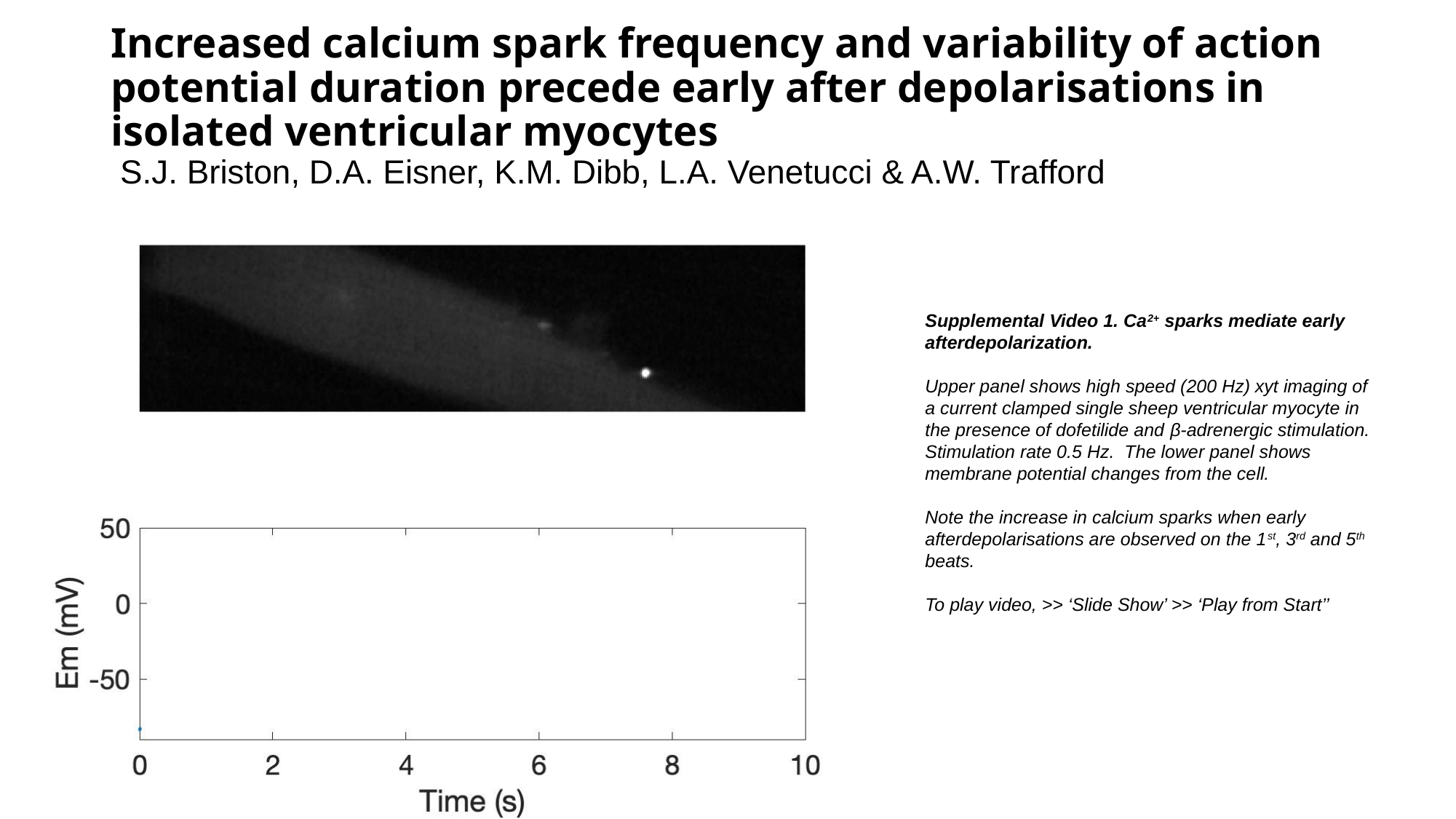

#### Increased calcium spark frequency and variability of action potential duration precede early after depolarisations in isolated ventricular myocytes  S.J. Briston, D.A. Eisner, K.M. Dibb, L.A. Venetucci & A.W. Trafford
Supplemental Video 1. Ca^2+^ sparks mediate early afterdepolarization.
Upper panel shows high speed (200 Hz) xyt imaging of a current clamped single sheep ventricular myocyte in the presence of dofetilide and β-adrenergic stimulation. Stimulation rate 0.5 Hz. The lower panel shows membrane potential changes from the cell.
Note the increase in calcium sparks when early afterdepolarisations are observed on the 1^st^, 3^rd^ and 5^th^ beats.
To play video, >> ‘Slide Show’ >> ‘Play from Start’’
